# Neuronal Culture Media Shape Protein Abundance and Turnover in Human iPSC-Derived Neurons

**DOI:** 10.64898/2026.11.25.769287

**Authors:** Jamison Shih, Ashley M. Frankenfield, Jiawei Ni, Hope Tracey, Ling Hao

## Abstract

Cultured neuron models are essential for studying neuronal biology and disease, yet it remains unclear how neuronal culture media shape the global proteome and protein turnover. Here, we compared human iPSC-derived neurons cultured in four different media: BrainPhys, DMEM/F-12, Neurobasal, or a 1:1 DMEM/F-12:Neurobasal mixture using label-free and dynamic SILAC proteomics. To enable protein turnover measurements in BrainPhys, we developed an in-house SILAC-compatible BrainPhys formulation that supported comparable viability and proteome coverage to commercial BrainPhys. BrainPhys produced a proteome and turnover landscape distinct from the other three media, which closely resembled one another. BrainPhys-cultured neurons showed lower relative abundance of DNA replication- and progenitor-associated proteins and higher relative abundance of synaptic and neuronal projection proteins, consistent with a more mature neuronal proteome. BrainPhys also globally prolonged protein half-lives, yielding a turnover profile more similar to later-stage primary neurons and *in vivo* brain. Despite their distinct basal proteomes, neurons cultured in different media exhibited a conserved proteomic response to acute lysosomal inhibition. Bafilomycin A1 treatment increased the relative abundance and synthesis of autophagy-associated proteins while decreasing the relative abundance of lysosomal hydrolases. Together, these findings identify neuronal culture medium as a critical determinant of neuronal proteome composition and dynamics and establish a SILAC-compatible BrainPhys formulation for future studies of neuronal proteostasis.

## Introduction

Cultured neuron models are essential for understanding neuron biology and modeling neurological diseases (1). Neuron cultures can be generated from primary animal tissues, embryonic tissues, or pluripotent stem cells (2, 3). Human induced pluripotent stem cell (iPSC)-derived neurons have become particularly valuable because they provide scalable human models and avoid species-specific differences that limit the translation of findings from animal systems (4–8). Furthermore, patient-derived iPSCs retain the genetic background associated with disease, enabling the study of pathogenic mechanisms and therapeutic responses in a clinically relevant context (1). However, cultured neurons are highly sensitive to experimental conditions. Factors including genetic stability, culture conditions, cellular heterogeneity, maturation state, and donor background can all contribute to molecular and functional variability (9–11). As proteomic approaches are increasingly applied to primary neurons, brain tissue, and iPSC-derived neuronal systems (12–18), understanding how controllable culture variables shape the neuronal proteome is essential for improving reproducibility and enabling meaningful comparisons across studies.

A key determinant of neuronal culture conditions is the culture medium itself. It provides the ionic environment, basic nutrients, hormones, and protein supplements that neurons need for survival, excitability, and synaptic function (19–22). Commonly used basal media for neuronal culture include DMEM/F-12 (hereafter DMEM), Neurobasal, a mixture of the two (DMEM+Neurobasal), and BrainPhys. Unlike conventional media that mainly support neuronal survival, BrainPhys was formulated to mimic the physiological composition of human cerebrospinal fluid and better support neuronal physiological function (23). Consistent with the initial discovery, subsequent studies also demonstrated that BrainPhys enhances neuronal electrical activity, synaptic transmission, and functional maturation across multiple neuronal models (24–26). Despite these well-characterized functional benefits, it remains unclear how different basal media influence the neuronal proteome on a proteome-wide scale. How culture medium composition alters proteostasis and specific molecular pathways has not been systematically investigated.

Neurons rely heavily on protein quality control systems to maintain homeostasis because of their postmitotic nature and highly polarized morphology (2). Neuronal proteostasis is sustained through a dynamic balance between protein synthesis and degradation, with protein degradation mediated primarily by the ubiquitin–proteasome system and the autophagy–lysosome pathway (27). Disruption of these proteostatic pathways leads to the accumulation of misfolded and aggregated proteins, a hallmark of aging and many neurodegenerative diseases (28–30). Protein turnover rate or half-life provides a quantitative measure of proteostasis by reflecting the net balance between protein synthesis and degradation (31–35). We and others have used dynamic Stable Isotope Labeling by Amino Acids in Cell Culture (dSILAC)-based quantitative proteomics to measure apparent protein half-lives in cultured neurons under both physiological and pathological conditions (16, 17, 36–42). However, these studies were all performed using conventional DMEM or Neurobasal media, because SILAC-compatible BrainPhys medium is not available as an off-the-shelf commercial product and must be custom manufactured in large batches at substantial cost. Consequently, it remains unknown how BrainPhys medium affects neuronal proteostasis and protein turnover.

To enable quantitative analysis of protein turnover in BrainPhys-cultured neurons, we developed a SILAC-compatible BrainPhys formulation based on the published defined composition (23) and verified its performance against commercial BrainPhys. We investigated how neuronal culture medium shapes protein abundance and turnover in human iPSC-derived neurons by comparing neurons maintained in BrainPhys, DMEM, Neurobasal, or DMEM+Neurobasal (1:1) using label-free and dSILAC proteomics analyses. We further examined whether culture medium influences neuronal responses to proteostasis stress by inhibiting lysosomal degradation with Bafilomycin A1 (BafA1) treatment (43). Together, our findings identify neuronal culture medium as a key determinant of steady-state proteome composition and protein turnover in human iPSC-derived neurons, while establishing a SILAC-compatible BrainPhys formulation for future studies of neuronal proteome dynamics.

## Experimental Procedures

### Experimental design and statistical rationale

To evaluate how neuronal media impact protein abundance and turnover, four commonly used neuronal media were compared. Three or four biological replicates (separate culture wells) from the same batch of differentiation were used for each medium in each type of experiment. Different types of the experiments (label-free, dSILAC, and drug treatment) were independently performed on different batches of differentiation on different days. Protein turnover measurement was conducted using a 4-day dSILAC time point as described previously with heavy-labeled medium containing ^13^C_6_,^15^N_2_ lysine (Lys8) and ^13^C_6_,^15^N_4_ arginine (Arg10) (17). For drug treatment with BafA1 vs. DMSO, both label-free and dSILAC experiments were conducted for 24 h to match the drug treatment time point. Differences in protein abundance and half-life were assessed using a moderated t-test as implemented in the limma package (44), with significance thresholds of |log_2_ FC| > 1 and Benjamini-Hochberg-adjusted p (p_adj_) < 0.05. Asterisks indicate adjusted p-values: *padj < 0.05, **padj < 0.01, ***padj < 0.001. Bar graphs show mean ± standard deviation (s.d.) with individual biological replicates shown as points.

### Human iPSC-neuron culture

Human iPSC-derived neurons were generated from the WTC11 iPSC line (Coriell GM25256, male) using the established i^3^Neuron technology, which stably expresses a doxycycline-inducible NGN2 cassette at the AAVS1 locus (45). Detailed protocol was described previously (46, 47). Briefly, to start differentiation, iPSCs were dissociated with Accutase (Corning) and replated at 5 million cells per 10-cm dish on Matrigel-coated dishes in 10 mL of N2 induction medium with DMEM/F12 supplemented with 1× N2 (Gibco), 1× MEM non-essential amino acids (0.1 mM; Gibco), 2 mM L-glutamine (Gibco), and 2 µg/mL doxycycline (Sigma). The medium was fully replaced daily, and ROCK inhibitor Y-27632 (10 µM) was added to the medium only for day 0 to prevent apoptosis. On day 3, cells were replated at 1 million cells per well on 12-well plates in 1 mL neuronal medium for each well. Plates were pre-coated with 0.1 mg/mL poly-L-ornithine (Sigma) in borate buffer (10 mM boric acid, 2.5 mM sodium tetraborate, 7.5 mM NaCl, pH 8.4) and washed with sterile water and air-dried. From day 3 to day 14, neuronal cultures were maintained in one of four neuronal media: BrainPhys (STEMCELL Technologies), DMEM/F12 (Gibco), Neurobasal (Gibco), or DMEM+Neurobasal (1:1) with medium changes every other day. All four media contained 1× B27 Plus (Gibco), 10 ng/mL each of BDNF, GDNF, and NT-3 (PeproTech), 0.2 µg/mL laminin (Gibco), and 2 µg/mL doxycycline from day 3 to day 14. DMEM/F12, Neurobasal, and DMEM/F12+Neurobasal (1:1) were additionally supplemented with 1× N2 and 0.1 mM non-essential amino acids. Neurobasal, which lacks L-glutamine, was supplemented with 2 mM L-glutamine. BrainPhys already contains 0.5 mM L-alanyl-L-glutamine, and DMEM/F12 contains 2.5 mM L-glutamine, so these media received no additional glutamine. No antibiotics were used at any stage. Final concentrations of all basal-medium components and supplements are listed in **Supplemental Table S1**. For BafA1 treatment, neurons were maintained in BrainPhys or DMEM/F-12 medium. On day 13, neurons were treated with 50 nM BafA1 (Cayman Chemical) or DMSO (0.1%) for 24 h before harvesting.

To measure protein turnover in neurons, dSILAC labeling was conducted in neurons cultured in four different media. Commercial SILAC media (lacking arginine and lysine) were used for DMEM, Neurobasal, and DMEM+Neurobasal (1:1) media. A homemade SILAC-compatible BrainPhys medium was created based on the published BrainPhys formulation (23). On day 10, neurons were gently washed twice with warm PBS and switched to heavy medium containing ^13^C ^15^N lysine (Lys8) and ^13^C ^15^N arginine (Arg10) (Cambridge Isotope). Neurons were maintained in heavy media for 4 days with warm half-media change on day 12 and harvested on day 14. For BafA1 treatment, heavy medium containing BafA1 was added to day 13 neurons for 24 h before harvesting.

### Proteomics sample preparation

Neurons were washed twice with warm PBS and harvested using ice-cold cell lysis buffer (8M Urea, 150 mM sodium chloride, 50 mM Tris-HCl buffer) and sonicated in a QSonica Q700 sonicator for 15 min using alternating 30 s on, 20 s off cycles in an ice-cold water bath. Protein concentrations were measured using the Bio-Rad detergent-compatible protein assay kit and normalized across all samples. Bottom-up proteomics were conducted as previously described (14, 48). Briefly, disulfide bonds were reduced using 5 mM TCEP for 40 min, and alkylated using 15 mM IAA for 30 min in the dark on a ThermoMixer shaking at 1,200 rpm at 37 °C. Samples were digested using 1:20 enzyme-to-protein ratio of Trypsin/Lys-C (Promega) for 16 h at 37 °C on the ThermoMixer and the reaction was quenched using 10% trifluoroacetic acid until the pH < 2. All samples were desalted using a Waters Oasis HLB plate, dried down using a SpeedVac, and stored at -30 °C. Label-free peptide samples and the four-media dSILAC comparison samples were processed without offline fractionation. For the BafA1 treatment experiment, dSILAC peptides were fractionated on a Waters XBridge BEH130 C18 column (3.5 μm, 130 Å, 4.6 mm × 250 mm) using a Vanquish HPLC, as previously described (17). Buffer A was 10 mM ammonium formate in water and buffer B was 10 mM ammonium formate in 90% acetonitrile, both adjusted to pH 10 with ammonium hydroxide. Fractions were collected with a Thermo Fraction Collector at 60 s intervals and a flow rate of 0.5 mL/min. The gradient held 5% B for 5 min, increased to 10% B over 2 min, reached 60% B by 35 min, held 90% B for 10 min, and re-equilibrated at 5% B for 10 min; fractions collected between 12 and 60 min were aggregated into six fractions per replicate, dried by SpeedVac, and stored at -30 °C.

### LC-MS/MS analysis

All peptide samples were dried and reconstituted in 0.1% formic acid (FA). LC-MS/MS analysis was performed on a Dionex Ultimate 3000 RSLC system coupled to a Thermo Fisher Q-Exactive HF-X mass spectrometer. Buffer A was 0.1% FA in water and buffer B was 0.1% FA in acetonitrile. Peptides were separated over a 182- or 272-min active gradient on an EASY-Spray PepMap RSLC C18 column (2 µm, 100 Å, 75 µm × 50 cm or 75 cm) held at 55 °C with a flow rate of 0.25 µL/min. Peptides from offline fractionation were separated over a 76 min active gradient on a 25 cm column. Data-independent acquisition (DIA) was conducted by performing a survey scan from *m/z* 400 to 1000 with a resolving power of 60K at *m/z* 200 full-width half-maximum (FWHM), automatic gain control (AGC) of 1 × 10^6^, and maximum injection time (MaxIT) of 60 ms. Precursors were fragmented using a normalized collision energy (NCE) of 30%. Fragments were detected using an *m/z* 8.0 staggered isolation pattern with 75 sequential DIA windows with a resolving power of 15K at *m/z* 200 FWHM, MaxIT of 40 ms, and an NCE of 30%.

### Proteomics data analysis

DIA proteomics data were analyzed using the directDIA workflow in Spectronaut software (49). The media-comparison datasets were processed with Spectronaut v19, and the BafA1 label-free and dSILAC datasets were processed with Spectronaut v20. Protein identification was achieved by searching against a reviewed Swiss-Prot *Homo sapiens* database (UniProtKB/Swiss-Prot release 2024_01, 20,433 entries, for the media-comparison datasets; release 2025_04, 20,420 entries, for the BafA1 datasets) and a custom neuron culture contaminant FASTA library (https://github.com/HaoGroup-ProtContLib) (48). Trypsin/P was used as the enzyme with a maximum of two missed cleavages. Carbamidomethylation of cysteine was set as a fixed modification, with methionine oxidation and protein N-terminal acetylation as variable modifications. Our recently optimized dSILAC Spectronaut data analysis parameters were used for dSILAC data (39). Briefly, Lys8/Arg10 was added as an additional label, *in-silico* generation of the missing isotopic channel with a labeled workflow was enabled, and only y-ions were used for quantification (50). Identifications were filtered at 1% FDR (precursor and experiment-wide protein q-value ≤ 0.01). Cross-run normalization was enabled for label-free abundance datasets and disabled for the dSILAC datasets.

All downstream data analyses were performed in R (v. 4.5.2) as described previously (17, 39). Heavy-to-light (H:L) ratios were computed at the peptide level, and outlier ratios outside 0.02-100 were excluded. For the media-comparison dSILAC dataset, peptide half-life was calculated using a first-order kinetics model, t_1/2_ = t_d_ ⋅ ln(2) / ln(1+H:L), where *t*_d_ represents the labeling time in days. Protein half-life was defined as the harmonic mean of all peptides belonging to each protein. For the BafA1 treatment dSILAC experiments, offline fractions from each replicate were combined after Spectronaut processing, aggregated to the peptide level. Heavy and light peptides were then analyzed separately for BafA1 vs. ctrl groups to account for non-steady state protein dynamics. Two-dimensional GO-enrichment analyses were computed based on the 2D-annotation enrichment framework (51). Principal component analysis was performed using the prcomp function in R. For label-free abundance data, missing values were not imputed; principal component analysis and hierarchical clustering used only proteins quantified in all samples, and each pair of conditions was compared separately using limma on log2-transformed intensities. GO over-representation analysis was performed using a hypergeometric test with Benjamini-Hochberg correction. Protein interaction networks were retrieved from the STRING database (v12.0; Homo sapiens; full network; combined score ≥ 0.4) (52). Custom code for analysis is available at https://github.com/HaoGroup. To assess precursor processing of lysosomal hydrolases, we quantified a tryptic peptide of CTSL that spans the UniProt-annotated propeptide-mature cleavage site (junction peptide), which can only arise from the unprocessed precursor. In each sample, the total protein abundance was estimated by Tukey’s median polish of log_2_ intensities of peptides from the mature region of the same protein, excluding junction and propeptide peptides (53).

### Fluorescence microscopy

Microscopy imaging was performed using a CELENA® S Digital Cell Imaging System inverted fluorescence microscope. Cell viability was assessed using the Invitrogen LIVE/DEAD green/red viability assay according to the manufacturer’s protocol. Viability was quantified from the raw green/live and red/dead fluorescence images. The same field selection criteria were used for all conditions and regions of cell detachment was avoided. Illumination intensity, exposure time, and gain were held constant across all wells and conditions (GFP channel: 5% light intensity, 30 ms exposure, 5 dB gain; RFP channel: 50% light intensity, 200 ms exposure, 5 dB gain). Each image was thresholded using the same MaxEntropy automatic threshold method in ImageJ. Pixels with intensity greater than or equal to the resulting threshold were counted as positive. Percent live viability was calculated as live-positive pixels / (live-positive pixels + dead-positive pixels) × 100. Microscopy data analysis was conducted using ImageJ (54).

### SDS-PAGE and immunoblotting

Neurons cultured in BrainPhys were treated with 25 nM BafA1 or 0.1% DMSO for 24 h (n = 3 wells per group) and prepared as described in the Proteomics sample preparation section. Equal amounts of protein (7 μg per lane) were mixed with Laemmli sample buffer (Bio-Rad) containing 100 mM DTT and denatured at 70 °C for 10 min. Proteins were separated on 4-12% Bis-Tris mini protein gels (1.0 mm; Invitrogen) and transferred onto nitrocellulose membranes using a Trans-Blot Turbo Transfer System (Bio-Rad). Membranes were blocked with 5% nonfat dry milk in TBST for 1 h at room temperature and incubated overnight at 4 °C with primary antibodies against cathepsin L (CTSL; Proteintech #10938-1-AP, 1:1000) and GAPDH (Invitrogen #AM4300, 1:5000), followed by HRP-conjugated secondary antibodies (Proteintech #SA00001-1, #SA00001-2) for 1 h at room temperature. Chemiluminescence was detected with enhanced chemiluminescence (ECL) substrate on a ChemiDoc imaging system (Bio-Rad). Band intensities were quantified using Image Lab software (Bio-Rad) and normalized to GAPDH. Pro-CTSL and mature CTSL band intensities were compared between groups using a two-tailed Welch’s t-test.

## Results

### BrainPhys shifts the steady-state proteome toward a less proliferative, more synaptic profile

To determine how neuronal culture media influence steady-state protein expression in human iPSC-derived neurons, we maintained NGN2-induced iPSC-derived neurons from day 3 to day 14 in four medium systems (BrainPhys, DMEM, Neurobasal, or DMEM+Neurobasal) and performed bottom-up proteomics on day 14 neurons (**Supplemental Table S2**). Neuron morphology and viability were comparable across all four culture conditions (**Fig. 1A**). Interestingly, total protein yield per million plated neurons was modestly lower in BrainPhys than in other media, which may suggest a less proliferative cellular state (**Fig. 1B**). Proteomic depths from equal protein inputs were comparable across all four conditions (**Fig. 1C, D, Supplemental Fig. S1**). Principal component analysis of the proteomic profiles revealed that BrainPhys produced a proteome distinct from those of the other three media, whereas DMEM, Neurobasal, and DMEM+Neurobasal clustered closely together (**Fig. 1E**). Pairwise correlation and hierarchical clustering further supported this separation (**Fig. 1F, Supplemental Fig. S1**).

**Fig. 1.**
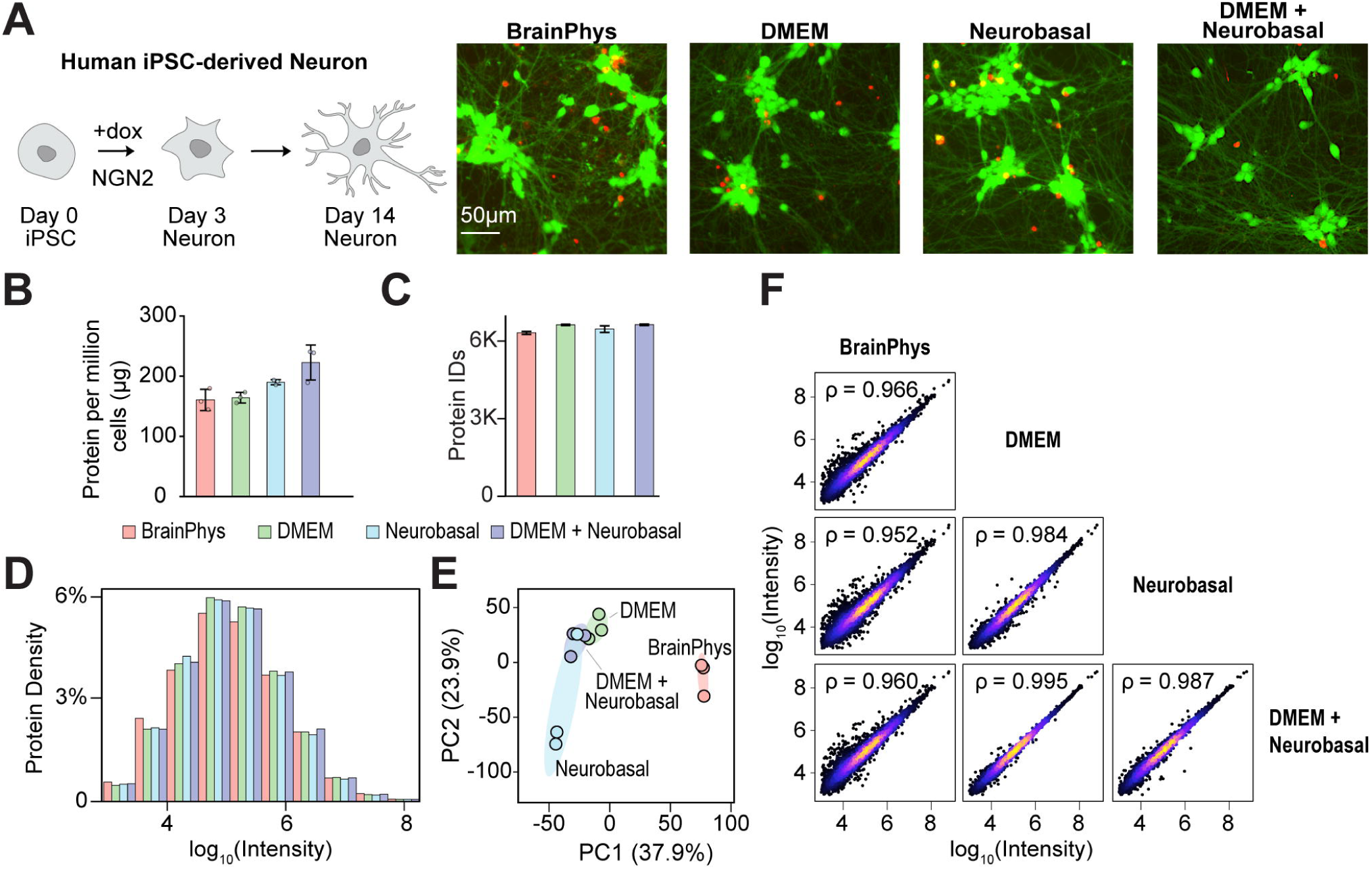
Comparison of human iPSC-derived neurons cultured in four neuronal media by microscopy and global proteomics. (A) Schematic of NGN2-induced iPSC-neuron differentiation and representative fluorescence images of day 14 neurons cultured in each medium. Live cells fluoresce green and dead cells fluoresce red in the LIVE/DEAD cell viability assay. (B) Total protein yield (μg) per million plated neurons (mean ± SD, n=3) (C) Numbers of quantified proteins (n = 3). (D) Density histograms of log_10_ protein intensities, averaged across biological replicates. (E) PCA of protein intensities. (F) Pairwise correlation of protein abundances with Spearman’s correlation coefficient (ρ).

Consistent with these global differences, differential abundance analysis comparing BrainPhys with each of the other media yielded highly similar volcano plot patterns (**Fig. 2A**). Across these comparisons, 350 proteins showed consistently lower relative abundance in BrainPhys, with significant enrichment for nuclear, DNA replication, and cell division pathways such as CMG and MCM complexes (**Fig. 2B**). In contrast, 62 proteins showed consistently higher relative abundance in BrainPhys and were enriched in synaptic and neuronal projection components (**Fig. 2C**). A heatmap of representative proteins illustrated this coordinated shift, with BrainPhys exhibiting lower relative abundance of DNA replication proteins (e.g., MCM, POLD, RFC, PCNA) and higher relative abundance of synaptic proteins (e.g., DLG4/PSD-95, CACNA1B, SYT3), whereas the proteomic profiles of DMEM, Neurobasal, and DMEM+Neurobasal remained highly similar (**Fig. 2D; Supplemental Fig. S2**). Persistent expression of DNA-replication and cell-cycle proteins in iPSC-neuron systems is often associated with residual progenitors or incompletely differentiated cells (11, 55, 56). Therefore, the lower relative abundance of these proteins together with the higher relative abundance of synaptic proteins on day 14 iNeurons suggests that BrainPhys promotes a less proliferative and more mature neuronal proteome.

**Fig. 2.**
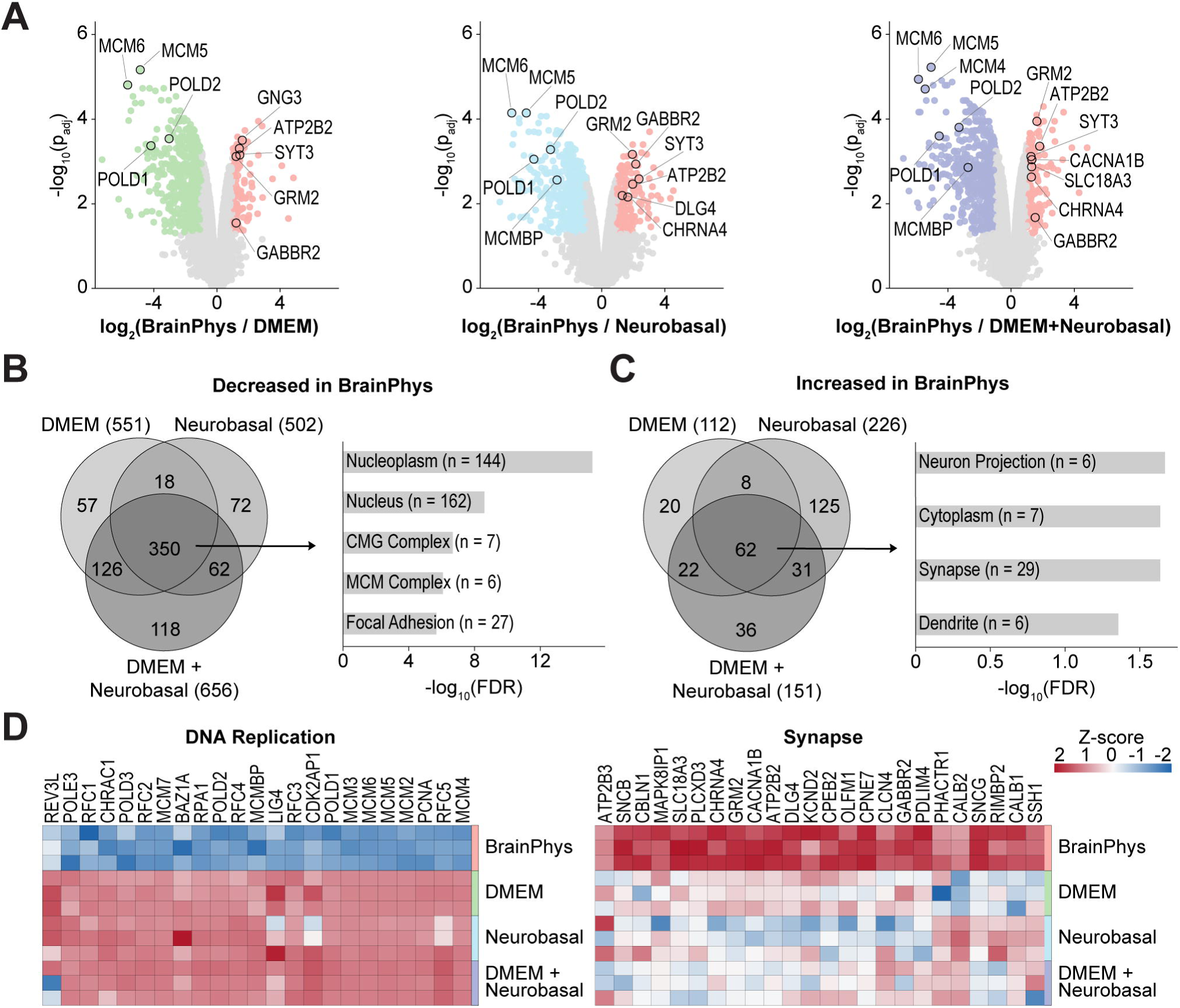
BrainPhys promotes a more synaptic and less proliferative neuronal proteome compared to other conventional media. (A) Volcano plots comparing relative protein abundance in BrainPhys with each of the other media (n=3). Differentially abundant proteins were defined by |log2FC| > 1 and adjusted p < 0.05. (B) Venn diagram showing the overlap of proteins consistently decreased in BrainPhys and the enriched GO terms. (C) Venn diagram showing the overlap of proteins consistently increased in BrainPhys and the enriched GO terms. (D) Heatmaps of representative proteins from the enriched DNA replication and synapse pathways.

### In-house formulated SILAC-compatible BrainPhys enables protein turnover measurement and broadly prolongs neuronal protein half-lives

To determine whether the BrainPhys-induced changes in steady-state proteome expression were accompanied by altered protein turnover, we performed dSILAC proteomics in neurons cultured in each of the four media. Because SILAC-compatible BrainPhys is not commercially available as an off-the-shelf product, we first established an in-house SILAC-compatible BrainPhys formulation based on the published defined composition (23) (**Fig. 3A, Supplemental Table S1**). This in-house formulated medium supported neuron morphology and viability comparable to those of commercial BrainPhys (**Fig. 3B**). Label-free proteomics further demonstrated a high correlation between neurons cultured in the in-house and commercial BrainPhys media (**Fig. 3C, D**). We then conducted dSILAC proteomics to measure apparent protein half-lives across the four media **(Fig. 3E, Supplemental Table S3)**. The number of quantified proteins/peptides as well as measurement reproducibility across replicates were comparable across all four media (**Supplemental Fig. S3A, B**).

**Fig. 3.**
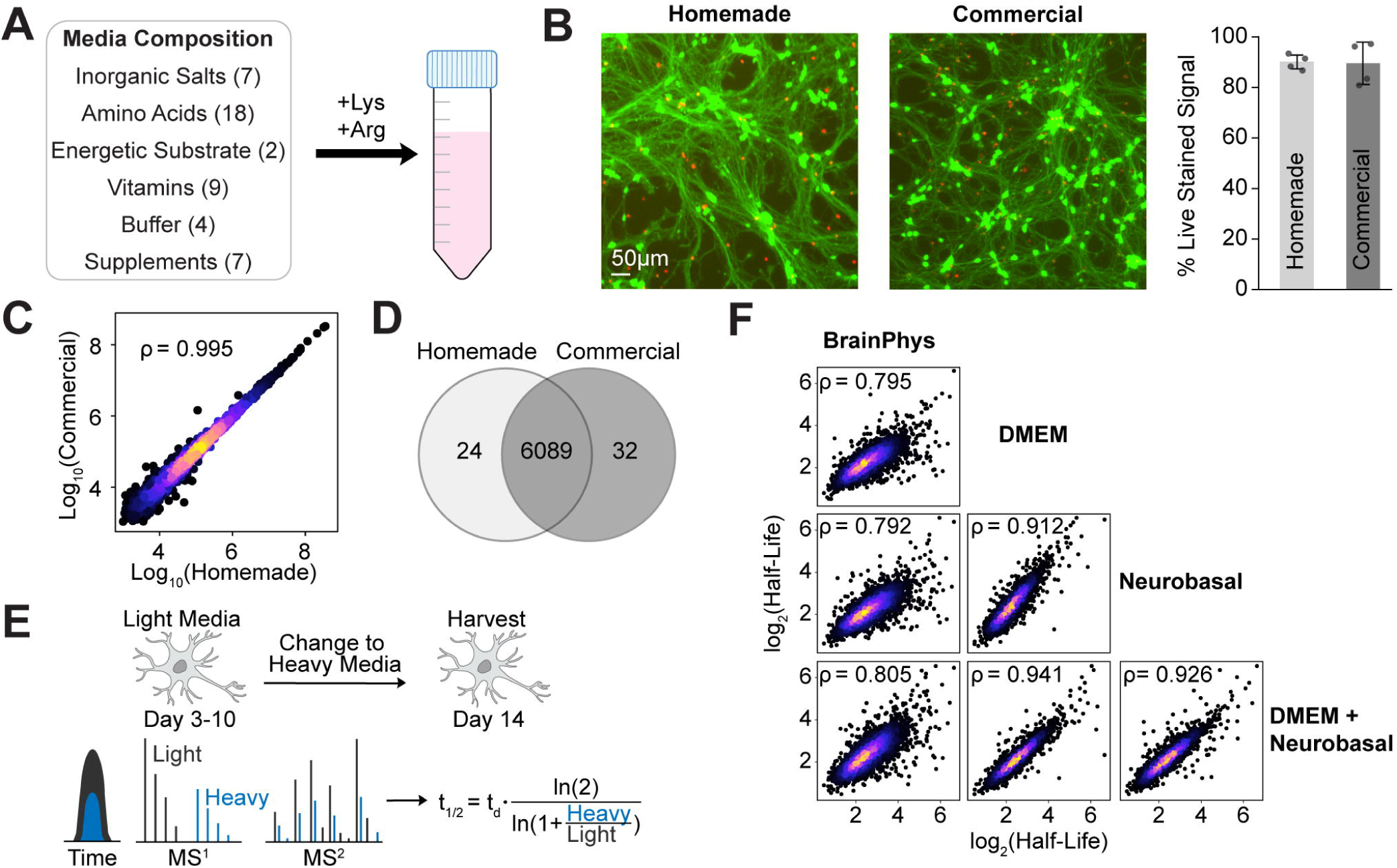
In-house formulated SILAC-compatible BrainPhys enables quantitative protein turnover measurements. (A) Composition of the in-house SILAC-compatible BrainPhys medium based on the published BrainPhys formulation. Numbers in parentheses indicate the number of components in each category. (B) Representative live/dead fluorescence images of day 14 neurons and viability comparison between in-house and commercial BrainPhys. (C) Pairwise correlation of protein abundances between in-house and commercial BrainPhys with Spearman’s ρ. (D) Venn diagram showing the overlap of quantified proteins. (E) Experimental workflow for dSILAC-based protein turnover measurements. Neurons were switched from light to heavy medium on day 10 and harvested on day 14; half-lives were calculated from heavy/light ratios using a first-order kinetic model with t_d_ = 4 days. (F) Pairwise correlation of protein half-lives across four neuronal media with Spearman’s ρ.

Despite similar analytical performance, BrainPhys exhibited a distinct global protein turnover profile. BrainPhys separated from the other three media by half-life correlation and hierarchical clustering (**Fig. 3F, Supplemental Fig. S3C**). PCA also showed distinct separation of BrainPhys from the other three media, with the greatest separation along PC1 (58.5%) (**Fig. 4A**). Volcano plots and half-life distribution consistently revealed widespread protein half-life prolongation in BrainPhys relative to each of the other media (**Fig. 4B, Supplemental Fig. S3C**). These findings indicate that BrainPhys broadly stabilizes the neuronal proteome rather than selectively altering turnover of only a small subset of proteins. To place these observations in the context of neuronal maturation, we compared our protein turnover measurement with published datasets from primary neurons and mouse brain tissues (13, 40, 57). Interestingly, BrainPhys clustered more closely with in vivo mouse cortex datasets, whereas DMEM, Neurobasal, and DMEM+Neurobasal grouped with embryonic primary neuron cultures (**Fig. 4C**). Together, these comparisons suggest that BrainPhys establishes a protein turnover landscape more closely resembling mature neuronal systems.

**Fig. 4.**
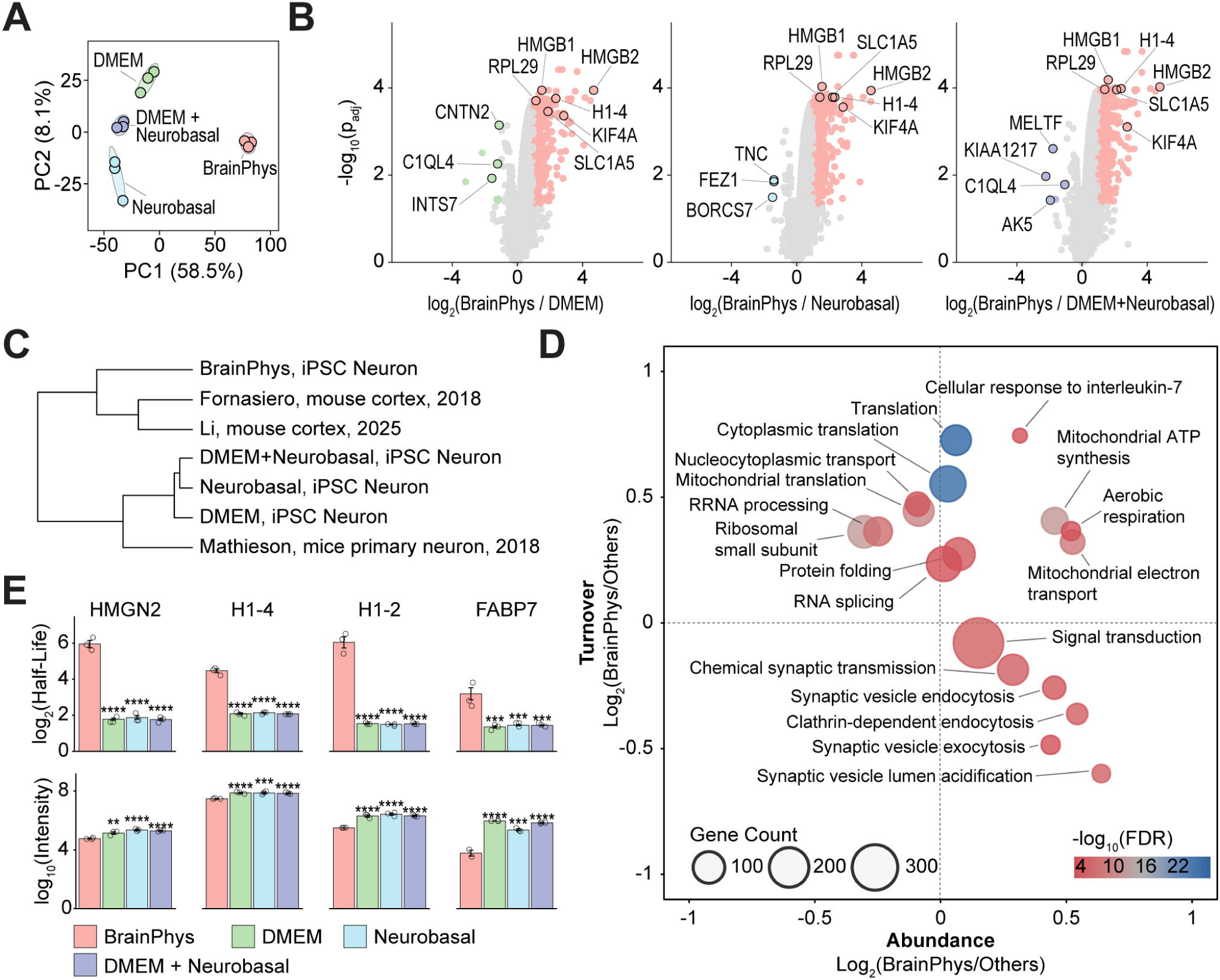
BrainPhys prolongs protein half-lives and promotes a mature proteostatic state. (A) PCA of protein half-lives (n = 3). (B) Volcano plots of half-life differences between BrainPhys and each other medium (n = 3). (C) Hierarchical clustering of replicate-averaged half-lives with published primary neuron and mouse brain datasets. (D) Two-dimensional GO biological process enrichment for abundance and half-life changes in BrainPhys versus the other media. Axes show log_2_(BrainPhys/others) for abundance (x) and half-life (y), using means from BrainPhys (n = 3) and the other media (n = 9). Circle size indicates proteins per term; color indicates −log_10_(FDR). (E) Half-lives (top) and relative abundances (bottom) of representative proteins (n = 3).

To identify the biological pathways underlying these global turnover differences, we integrated protein abundance and turnover measurements across the four media. Two-dimensional GO-enrichment analysis showed that proteins involved in translation, RNA-processing, and mitochondrial processes shifted toward longer half-lives in BrainPhys, whereas synaptic proteins shifted toward higher relative abundance and shorter half-lives, suggesting active remodeling of synaptic machinery (**Fig. 4D**). ATP-synthesis, aerobic-respiration, and electron-transport terms shifted toward higher relative abundance and longer half-lives in BrainPhys. At the individual-protein level, joint abundance-turnover analysis identified proteins that were lower in relative abundance but longer-lived in BrainPhys, predominantly nuclear and chromatin-associated proteins, including linker histones, HMG proteins, and DNA replication factors, consistent with a more postmitotic nuclear proteome (**Fig. 4E**). Collectively, these results demonstrate that BrainPhys reshapes both protein abundance and turnover in pathways associated with neuronal maturation, metabolic specialization, and establishment of a mature proteostatic state.

### Different neuronal media exhibit a conserved protein abundance response to lysosomal inhibition

Having established that BrainPhys reshapes the steady-state neuronal proteome and protein turnover, we next asked whether neuronal culture medium also influences the proteomic response to proteostasis stress. We treated neurons with the lysosomal inhibitor BafA1 and investigated changes in both protein abundance and turnover (**Fig. 5A**). Because DMEM, Neurobasal, and DMEM+Neurobasal exhibited highly similar steady-state protein abundance and turnover profiles, we selected DMEM as a representative conventional medium for comparison with BrainPhys. The numbers of quantified proteins were comparable across media and treatment groups in label-free proteomics, but fewer H/L protein ratios were quantified in BrainPhys likely due to slower turnover profile (**Fig. 5B**). The PCA plot showed tight clustering of biological replicates and clear separation by both culture media and BafA1 treatment (**Fig. 5C**). Despite the distinct steady-state proteomes of BrainPhys- and DMEM-cultured neurons, BafA1 induced remarkably similar changes in protein abundance in both media (**Fig. 5D**). A shared set of 47 proteins increased in relative abundance following BafA1 treatment, including autophagy receptors and adaptors (SQSTM1/p62, NBR1, TAX1BP1, and NCOA4), autophagic cargo and membrane-associated proteins, and the lysosomal biogenesis transcription factor TFE3 (**Fig. 5E, Supplemental Table S4**). GO enrichment analysis of these commonly increased proteins identified autophagosome- and synapse-related pathways (**Fig. 5E; Supplemental Fig. S4**). Although fewer proteins were significantly decreased in BrainPhys than in DMEM, both media shared reduced relative abundance of six soluble lysosomal hydrolases (**Fig. 5F**). Together, these results indicate that acute lysosomal inhibition elicits a largely conserved proteomic response in different neuronal media.

**Fig. 5.**
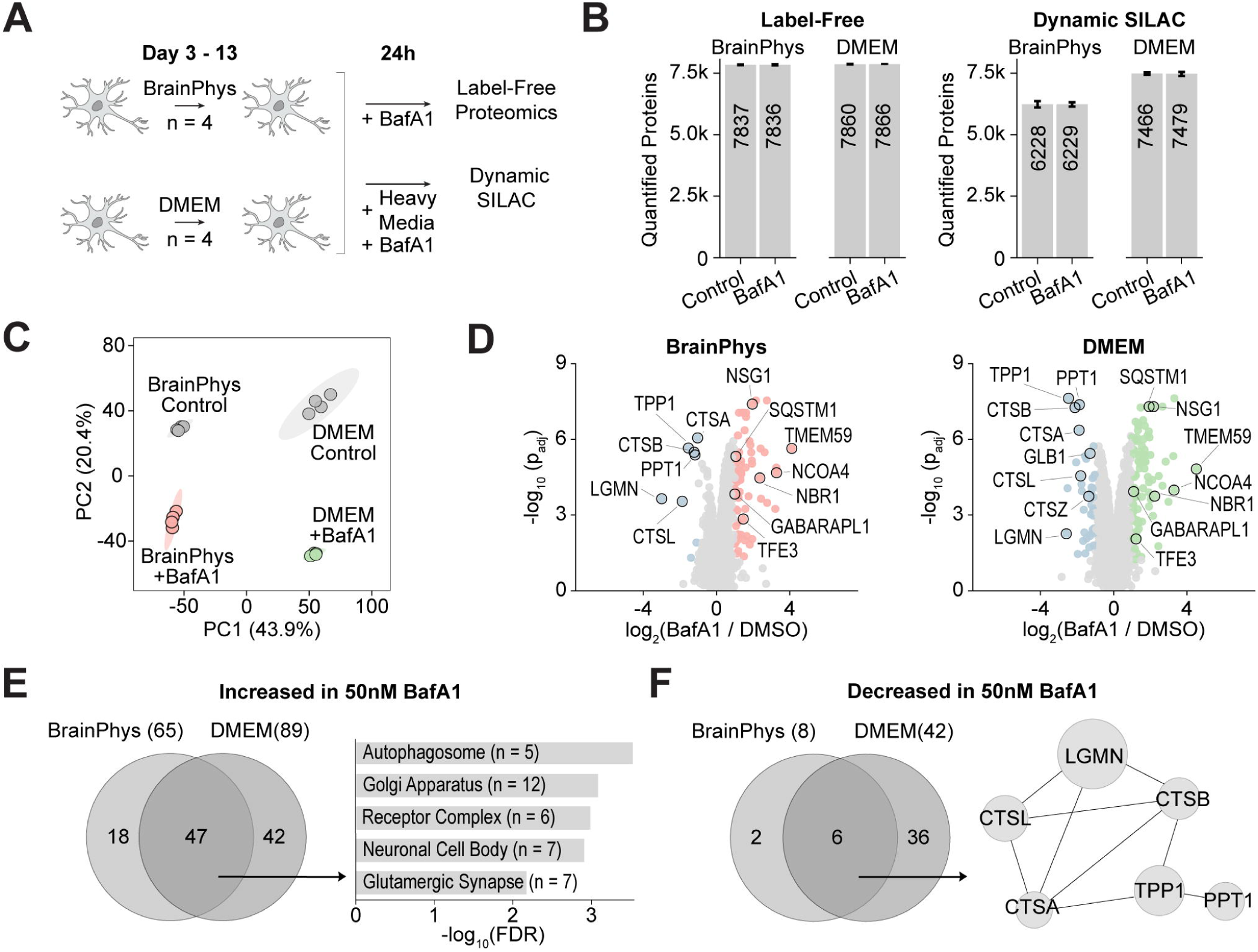
Global proteomic analysis of BrainPhys- and DMEM-cultured neurons treated with bafilomycin A1. (A) Experimental design for 24-h treatment with 50 nM BafA1 followed by label-free and dSILAC proteomics with 0.1% DMSO as the control (n = 4). (B) Numbers of quantified proteins in the label-free (left) and dSILAC (right) proteomics experiments (n = 4). (C) PCA of protein abundances across medium and treatment groups. (D) Volcano plots comparing BafA1-treated and control neurons in BrainPhys (left) and DMEM (right) (n = 4). (E) Venn diagram showing the overlap of proteins consistently increased following BafA1 treatment and the enriched GO terms (cellular component). (F) Venn diagram showing the overlap of proteins consistently decreased following BafA1 treatment. Node size is scaled to the mean absolute log_2_FC across BrainPhys and DMEM.

### Conserved proteomic responses to lysosomal inhibition reflect distinct effects on protein synthesis and clearance

Conventional protein half-life estimation assumes that the proteome is at steady state. Because acute BafA1 treatment blocks lysosomal degradation and disrupts this steady state, protein half-lives can no longer be reliably estimated under these conditions (31, 58, 59). Instead, we analyzed newly synthesized (heavy) and pre-existing (light) peptide pools separately to distinguish synthesis- and clearance-associated proteomic changes. BrainPhys showed fewer protein changes overall than DMEM, but both media displayed remarkably similar patterns of heavy- and light-protein changes (**Fig. 6A, B, Supplemental Fig. S5**). Proteins that increased in relative abundance following BafA1 treatment, including SQSTM1, GABARAPL1, NSG1/2, and SYT11, showed the largest increases in the newly synthesized (heavy) protein pool, whereas relatively little change was observed in the pre-existing (light) pool (**Fig. 6C**). This pattern suggests that the increased relative abundance of these canonical autophagy substrates reflects continued protein synthesis combined with impaired lysosomal clearance. In contrast, several soluble lysosomal hydrolases, including CTSL, TPP1, PPT1, and GLB1, decreased consistently across total, heavy, and light protein measurements in both BrainPhys and DMEM.

**Fig. 6.**
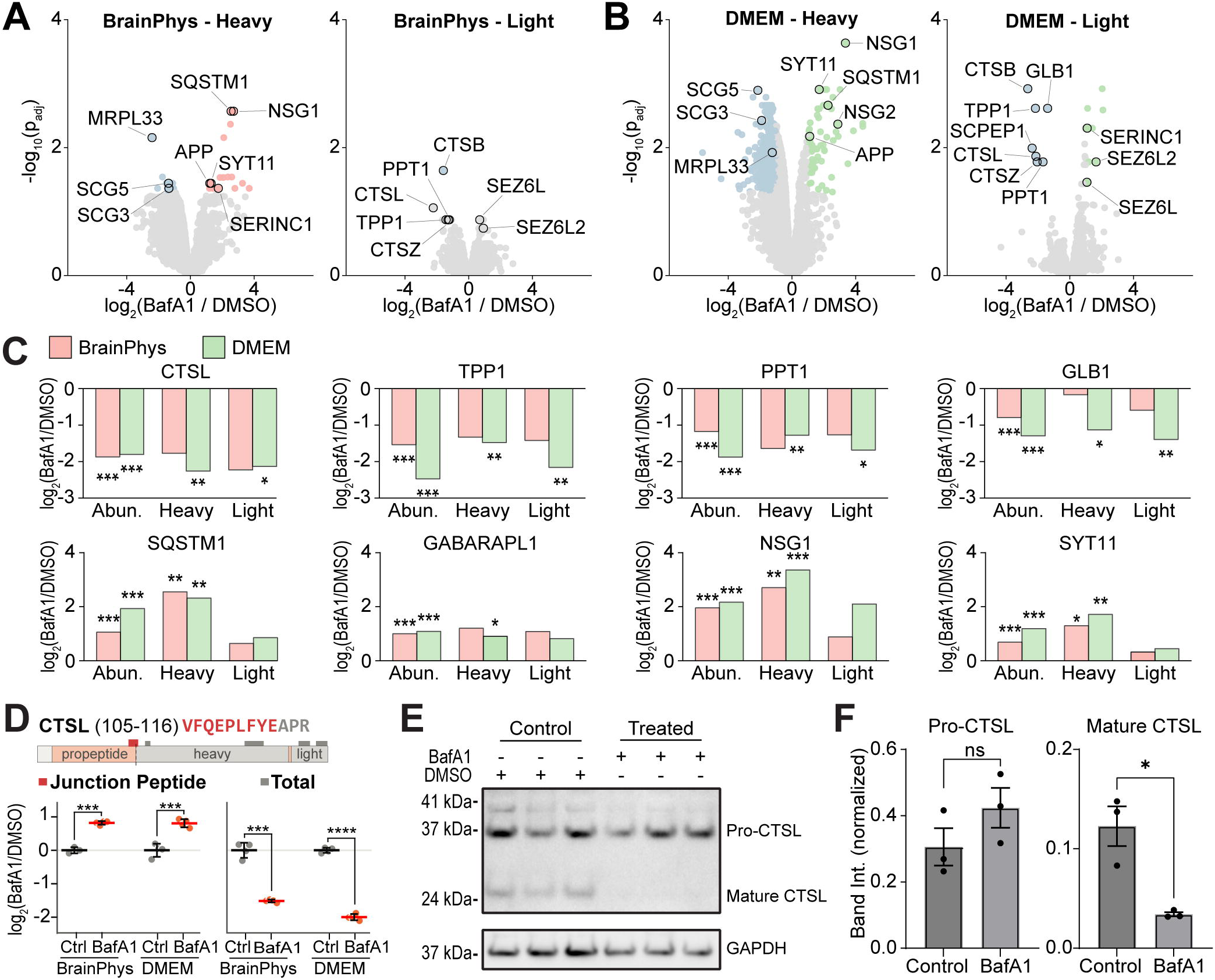
dSILAC analysis of newly synthesized and pre-existing proteins following bafilomycin A1 treatment. (A) Volcano plots comparing newly synthesized (heavy) and pre-existing (light) proteins following BafA1 treatment in BrainPhys medium (n = 4). (B) Volcano plots comparing newly synthesized and pre-existing proteins in DMEM medium (n=4). (C) Changes in total (abundance), newly synthesized (heavy), and pre-existing (light) protein pools for representative proteins following BafA1 treatment. (D) Abundance of CTSL peptides showing the junction peptide specific to pro-CTSL and mature-region peptides used to estimate total CTSL abundance (n=4). The junction peptide and total CTSL were each compared between BafA1 and DMSO by two-tailed Welch’s t-test. (E) Immunoblot of CTSL in BafA1-treated and control neurons showing pro-CTSL and mature CTSL levels (n=3). (F) Quantification of pro-CTSL and mature CTSL immunoblot bands with two-tailed Welch’s t-test.

Because lysosomal hydrolases undergo proteolytic processing to generate their mature forms, we next investigated whether the observed decrease in hydrolase abundance was associated with impaired protein maturation. We focused on cathepsin L (CTSL), one of the most significantly decreased hydrolases following BafA1 treatment, which undergoes well-characterized processing from its precursor to mature form. A CTSL peptide spanning the propeptide–heavy chain junction (VFQEPLFYEAPR, residues 105–116), specific to the unprocessed precursor, increased 1.8-fold following BafA1 treatment in both media. In contrast, CTSL abundance estimated from peptides within the mature region decreased 2.9-fold in BrainPhys and 4.0-fold in DMEM (**Fig. 6D**). Consistent with these proteomics results, immunoblotting confirmed a significant reduction in the mature CTSL heavy chain (∼24 kDa), accompanied by an increase in pro-CTSL (**Fig. 6E, F**). These findings indicate that the apparent decrease in CTSL abundance is associated with impaired proteolytic maturation and accumulation of its precursor rather than a uniform reduction in all CTSL forms. Together, these results reveal distinct responses to lysosomal inhibition, characterized by increased abundance of newly synthesized autophagy-associated proteins and reduced levels of mature lysosomal hydrolases, with largely conserved patterns across both neuronal culture media.

## Discussion

Although neuronal media are known to affect neuronal function, their global effects on protein abundance and turnover are largely unknown (23, 24). Here, we found that neuronal culture medium is a major determinant of protein abundance and turnover in human iPSC-derived neurons. Among the four media tested, BrainPhys produced a proteomic state distinct from DMEM, Neurobasal, and DMEM+Neurobasal, which were comparatively similar. BrainPhys-cultured neurons showed lower relative abundance of DNA-replication and cell-cycle proteins, including MCM, POLD, RFC, and PCNA proteins, together with higher relative abundance of synaptic and neuronal-projection proteins such as DLG4/PSD-95, CACNA1B, and SYT3 (11, 55, 56). These coordinated differences are consistent with a less proliferative and more synaptically differentiated proteomic state.

BrainPhys also broadly altered neuronal protein turnover. Protein half-lives were generally longer in BrainPhys than in the three conventional media, and the resulting turnover profile clustered more closely with later-stage primary-neuron and in vivo brain datasets (13, 40, 57). Nuclear and chromatin-associated proteins, including linker histones, HMG proteins, and DNA-replication factors, showed lower relative abundance but longer half-lives in BrainPhys, consistent with increased stability of nuclear structures in postmitotic cells (60, 61). In contrast, synaptic proteins tended to combine higher relative abundance with shorter relative half-lives, suggesting continued remodeling of synaptic machinery (62–64). Mitochondrial respiration and ATP-synthesis pathways also shifted toward higher relative abundance and longer half-lives, in agreement with the metabolic specialization associated with neuronal differentiation (65). Thus, BrainPhys appears to establish a proteostatic state that more closely resembles mature neuronal systems while maintaining active turnover of functionally dynamic proteins.

Despite these baseline differences, the acute response to lysosomal inhibition was largely conserved between different neuronal media. BafA1 blocks proton transport into the lysosomal lumen, with a resulting loss of acidification that impairs both autophagosome-lysosome fusion and luminal degradation (43, 66). BafA1 increased autophagy receptors and adaptors, including SQSTM1, NBR1, TAX1BP1, and GABARAPL1, as well as the lysosomal-biogenesis factor TFE3 (67). Separate analysis of the heavy- and light-protein pools indicated that the increased relative abundance of several autophagy-associated proteins was driven primarily by newly synthesized protein, consistent with continued synthesis combined with impaired lysosomal clearance. In contrast, soluble lysosomal hydrolases, including CTSL, TPP1, PPT1, and GLB1, decreased in total, newly synthesized, and pre-existing pools (67, 68). Junction-peptide analysis and immunoblotting of CTSL further indicated that BafA1 impaired conversion of the precursor to the mature enzyme, consistent with the requirement of an acidic lysosomal lumen for cathepsin activation. The lower relative abundance of hydrolases may therefore reflect impaired processing or stability of lysosomal enzymes following loss of luminal acidification. The parallel induction of autophagy receptors and TFE3 is also consistent with activation of a compensatory Coordinated Lysosomal Expression and Regulation (CLEAR)-network response (69). Overall, BafA1 triggered a largely conserved lysosomal-stress response across media, combining compensatory induction of autophagy-related proteins with loss of soluble lysosomal hydrolases.

In conclusion, neuronal medium composition should be treated as an important experimental variable in proteomic studies of cultured neurons. BrainPhys shifts both protein abundance and turnover toward a state more consistent with synaptically differentiated and later-stage neuronal systems, whereas the response to acute lysosomal inhibition remains broadly conserved across media. The SILAC-compatible BrainPhys formulation developed here enables future studies of neuronal protein dynamics under culture conditions designed to better support neuronal physiology.

## Acknowledgements

This study is supported by the NIH [R01NS121608] and Target ALS Basic Biology Consortia Grant [BB2026C4L2]. A.M.F. acknowledges the Willard & Marilynn Sweetser Scholarship from the Metropolitan Chapter of Achievement Rewards for College Scientists (ARCS) Foundation. The authors thank the Jiou Wang group from Johns Hopkins University for gifting the SILAC Neurobasal medium used in the study. The authors thank all Hao Lab members for providing feedback on the manuscript and their support.

## Data availability

All LC-MS/MS data have been deposited to the ProteomeXchange Consortium (70) via the PRIDE (71) partner repository with the dataset identifier PXD081410.

## Author contributions

A.M.F., J.N., L.H. Conceptualization; J.S., H.T. Formal analysis; J.S., A.M.F., J.N., H.T. Investigation; L.H. Resources; J.S. Writing - original draft; J.S., A.M.F., J.N., H.T., L.H. Writing - review & editing; L.H. Supervision; L.H. Project administration; L.H. Funding acquisition.

## Declaration of competing interests

The authors declare no competing interests. The authors have no financial relationship with STEMCELL Technologies, the manufacturer of BrainPhys.

## Abbreviations

AGC: automatic gain control
BafA1: bafilomycin A1
CLEAR: coordinated lysosomal expression and regulation
DIA: data-independent acquisition
DMEM: DMEM/F12 (Dulbecco’s Modified Eagle Medium/Nutrient Mixture F-12)
DMSO: dimethyl sulfoxide
dSILAC: dynamic stable isotope labeling by amino acids in cell culture
FA: formic acid
FDR: false discovery rate
FWHM: full-width half-maximum
GO: gene ontology
H:L: heavy-to-light ratio
IAA: iodoacetamide
iPSC: induced pluripotent stem cell
LFQ: label-free quantification
MaxIT: maximum injection time
NCE: normalized collision energy
NGN2: neurogenin 2
PBS: phosphate-buffered saline
PCA: principal component analysis
SILAC: stable isotope labeling by amino acids in cell culture
TCEP: tris(2-carboxyethyl)phosphine
t½: half-life

## Notes

### Competing Interest Statement

The authors have declared no competing interest.

## References

1. Dolmetsch, R., and Geschwind, D. H. (2011) The Human Brain in a Dish: The Promise of iPSC-Derived Neurons. Cell 145, 831–834

2. Levitan, I. B., and Kaczmarek, L. K. (2015) The Neuron (Oxford University Press)

3. Gordon, J., Amini, S., and White, M. K. (2013) in Neuronal Cell Culture. Methods in Molecular Biology (Humana Press, Totowa, NJ.), pp 1–8.

4. Hodge, R. D., Bakken, T. E., Miller, J. A., Smith, K. A., Barkan, E. R., Graybuck, L. T., Close, J. L., Long, B., Johansen, N., Penn, O., Yao, Z., Eggermont, J., Höllt, T., Levi, B. P., Shehata, S. I., Aevermann, B., Beller, A., Bertagnolli, D., Brouner, K., Casper, T., Cobbs, C., Dalley, R., Dee, N., Ding, S.-L., Ellenbogen, R. G., Fong, O., Garren, E., Goldy, J., Gwinn, R. P., Hirschstein, D., Keene, C. D., Keshk, M., Ko, A. L., Lathia, K., Mahfouz, A., Maltzer, Z., McGraw, M., Nguyen, T. N., Nyhus, J., Ojemann, J. G., Oldre, A., Parry, S., Reynolds, S., Rimorin, C., Shapovalova, N. V, Somasundaram, S., Szafer, A., Thomsen, E. R., Tieu, M., Quon, G., Scheuermann, R. H., Yuste, R., Sunkin, S. M., Lelieveldt, B., Feng, D., Ng, L., Bernard, A., Hawrylycz, M., Phillips, J. W., Tasic, B., Zeng, H., Jones, A. R., Koch, C., and Lein, E. S. (2019) Conserved cell types with divergent features in human versus mouse cortex. Nature 573, 61–68

5. Mertens, J., Marchetto, M. C., Bardy, C., and Gage, F. H. (2016) Evaluating cell reprogramming, differentiation and conversion technologies in neuroscience. Nat. Rev. Neurosci. 17, 424–437

6. Takahashi, K., Tanabe, K., Ohnuki, M., Narita, M., Ichisaka, T., Tomoda, K., and Yamanaka, S. (2007) Induction of Pluripotent Stem Cells from Adult Human Fibroblasts by Defined Factors. Cell 131, 861–872

7. Lu, J., Zhong, X., Liu, H., Hao, L., Huang, C. T. L., Sherafat, M. A., Jones, J., Ayala, M., Li, L., and Zhang, S. C. (2016) Generation of serotonin neurons from human pluripotent stem cells. Nat. Biotechnol. 34, 89–94

8. Collier, T. S., Sarkar, P., Rao, B., and Muddiman, D. C. (2010) Quantitative top-down proteomics of SILAC labeled human embryonic stem cells. J. Am. Soc. Mass Spectrom. 21, 879–889

9. Volpato, V., and Webber, C. (2020) Addressing variability in iPSC-derived models of human disease: guidelines to promote reproducibility. Dis. Model. Mech. 13,

10. Muñoz-Estrada, J., Mostafania, A., Halwatura, L., Haghani, A., Jiang, Y., and Meyer, J. G. (2026) Optimizing NGN2 Dosage Enhances the Neuronal Enrichment of iPSC-Derived Neuronal Cultures. Molecular & Cellular Proteomics 25, 101604

11. Shan, X., Zhang, A., Rezzonico, M. G., Tsai, M.-C., Sanchez-Priego, C., Zhang, Y., Chen, M. B., Choi, M., Andrade López, J. M., Phu, L., Cramer, A. L., Zhang, Q., Pattison, J. M., Rose, C. M., Hoogenraad, C. C., and Jeong, C. G. (2024) Fully defined NGN2 neuron protocol reveals diverse signatures of neuronal maturation. Cell Reports Methods 4, 100858

12. Sharma, K., Schmitt, S., Bergner, C. G., Tyanova, S., Kannaiyan, N., Manrique-Hoyos, N., Kongi, K., Cantuti, L., Hanisch, U.-K., Philips, M.-A., Rossner, M. J., Mann, M., and Simons, M. (2015) Cell type– and brain region–resolved mouse brain proteome. Nat. Neurosci. 18, 1819–1831

13. Li, W., Dasgupta, A., Yang, K., Wang, S., Hemandhar-Kumar, N., Chepyala, S. R., Yarbro, J. M., Hu, Z., Salovska, B., Fornasiero, E. F., Peng, J., and Liu, Y. (2025) Turnover atlas of proteome and phosphoproteome across mouse tissues and brain regions. Cell 188, 2267–2287.e21

14. Lee, G. Bin, Mazli, W. N. A. B., and Hao, L. (2024) Multiomics Evaluation of Human iPSCs and iPSC-Derived Neurons. J. Proteome Res. 23, 3149–3160

15. Shrestha, H. K., Sun, H., Yarbro, J. M., Lee, D., Liu, D., Wang, E., McReynolds, M., Zhang, N., Xie, B., Yang, S., Yu, K., Poudel, S., Li, Y., Yuan, Z.-F., Kong, D., Wang, M., Wang, Z., Niu, M., Wang, H., Zaman, M., Wang, J., Vanderwall, D. R., Sun, Y., Wu, Z., Chen, P.-C., Bai, B., High, A. A., Faura, J., Liu, C., Bennett, D. A., Johnson, E. C. B., Seyfried, N. T., Levey, A. I., Haroutunian, V., Serrano, G. E., Beach, T. G., DeTure, M., Kanekiyo, T., Petersen, R. C., Bu, G., McLean, P. J., Dickson, D. W., Rademakers, R., Yu, G., Wang, X., Zhang, B., and Peng, J. (2026) Pan-neurodegeneration proteomics reveals disease subtypes and molecular signatures. Cell 189, 3124–3143.e15

16. Hasan, S., Fernandopulle, M. S., Humble, S. W., Frankenfield, A. M., Li, H., Prestil, R., Johnson, K. R., Ryan, B. J., Wade-Martins, R., Ward, M. E., and Hao, L. (2023) Multi-modal proteomic characterization of lysosomal function and proteostasis in progranulin-deficient neurons. Mol. Neurodegener. 18, 87

17. Frankenfield, A., Ni, J., Zhang, T., Shih, J., Mazli, A., Li, W., Lo, E., Liu, Y., Wang, J., and Hao, L. (2026) A Deep Quantitative Proteome Turnover Platform for Human iPSC-derived Neurons.” Analytical Chemistry.

18. Lee, J., Chin, N., Zou, J., Mazli, W. N. A. B., Jarnik, M., Saidi, L., Xu, Y., Jeong, E., Suh, J., Replogle, J., Ward, M. E., Bonifacino, J. S., Zheng, W., Hao, L., and Ye, Y. (2025) CHIP protects lysosomes from CLN4 mutant-induced membrane damage. Nat. Cell Biol. 27, 1465–1481

19. Eagle, H. (1955) Nutrition Needs of Mammalian Cells in Tissue Culture. Science (1979). 122, 501–504

20. Dulbecco, R., and Freeman, G. (1959) Plaque production by the polyoma virus. Virology 8, 396–397

21. Brewer, G. J., Torricelli, J. R., Evege, E. K., and Price, P. J. (1993) Optimized survival of hippocampal neurons in B27-supplemented neurobasal^TM^, a new serum-free medium combination. J. Neurosci. Res. 35, 567–576

22. Sünwoldt, J., Bosche, B., Meisel, A., and Mergenthaler, P. (2017) Neuronal Culture Microenvironments Determine Preferences in Bioenergetic Pathway Use. Front. Mol. Neurosci. 10,

23. Bardy, C., Van Den Hurk, M., Eames, T., Marchand, C., Hernandez, R. V, Kellogg, M., Gorris, M., Galet, B., Palomares, V., Brown, J., Bang, A. G., Mertens, J., Böhnke, L., Boyer, L., Simon, S., and Gage, F. H. (2015) Neuronal medium that supports basic synaptic functions and activity of human neurons in vitro. Proceedings of the National Academy of Sciences 112,

24. Satir, T. M., Nazir, F. H., Vizlin-Hodzic, D., Hardselius, E., Blennow, K., Wray, S., Zetterberg, H., Agholme, L., and Bergström, P. (2020) Accelerated neuronal and synaptic maturation by BrainPhys medium increases Aβ secretion and alters Aβ peptide ratios from iPSC-derived cortical neurons. Sci. Rep. 10, 601

25. Zabolocki, M., McCormack, K., Van Den Hurk, M., Milky, B., Shoubridge, A. P., Adams, R., Tran, J., Mahadevan-Jansen, A., Reineck, P., Thomas, J., Hutchinson, M. R., Mak, C. K. H., Añonuevo, A., Chew, L. H., Hirst, A. J., Lee, V. M., Knock, E., and Bardy, C. (2020) BrainPhys neuronal medium optimized for imaging and optogenetics in vitro. Nat. Commun. 11, 5550

26. Jackson, T. C., Kotermanski, S. E., Jackson, E. K., and Kochanek, P. M. (2018) BrainPhys® increases neurofilament levels in CNS cultures, and facilitates investigation of axonal damage after a mechanical stretch-injury in vitro. Exp. Neurol. 300, 232–246

27. Kurtishi, A., Rosen, B., Patil, K. S., Alves, G. W., and Møller, S. G. (2019) Cellular Proteostasis in Neurodegeneration. Mol. Neurobiol. 56, 3676–3689

28. Hipp, M. S., Kasturi, P., and Hartl, F. U. (2019) The proteostasis network and its decline in ageing. Nat. Rev. Mol. Cell Biol. 20, 421–435

29. Ross, C. A., and Poirier, M. A. (2004) Protein aggregation and neurodegenerative disease. Nat. Med. 10, S10–S17

30. Yarbro, J. M., Han, X., Dasgupta, A., Yang, K., Liu, D., Shrestha, H. K., Zaman, M., Wang, Z., Yu, K., Lee, D. G., Vanderwall, D., Niu, M., Sun, H., Xie, B., Chen, P.-C., Jiao, Y., Zhang, X., Wu, Z., Chepyala, S. R., Fu, Y., Li, Y., Yuan, Z.-F., Wang, X., Poudel, S., Vagnerova, B., He, Q., Tang, A., Ronaldson, P. T., Chang, R., Yu, G., Liu, Y., and Peng, J. (2025) Human and mouse proteomics reveals the shared pathways in Alzheimer’s disease and delayed protein turnover in the amyloidome. Nat. Commun. 16, 1533

31. Pratt, J. M., Petty, J., Riba-Garcia, I., Robertson, D. H. L., Gaskell, S. J., Oliver, S. G., and Beynon, R. J. (2002) Dynamics of protein turnover, a missing dimension in proteomics. Mol. Cell. Proteomics 1, 579–591

32. Basisty, N., Meyer, J. G., and Schilling, B. (2018) Protein Turnover in Aging and Longevity. Proteomics 18,

33. Basisty, N., Holtz, A., and Schilling, B. (2020) Accumulation of “Old Proteins” and the Critical Need for MS-based Protein Turnover Measurements in Aging and Longevity. Proteomics 20, 1–8

34. Pino, L. K., Baeza, J., Lauman, R., Schilling, B., and Garcia, B. A. (2021) Improved SILAC Quantification with Data-Independent Acquisition to Investigate Bortezomib-Induced Protein Degradation. J. Proteome Res. 20, 1918–1927

35. Saha, S., Verma, R., and Nita-Lazar, A. (2026) Kinetic control of macrophage death by PTM-turnover crosstalk in infection and inflammation. Cell Chem. Biol. 33, 1038–1057

36. Liu, Y., Huang, Z., Hsu, Y.-W., Deme, P., Frankenfield, A. M., Wu, S., Zhao, X., Liu, H., Zhang, T., Alexander, E. J., Liu, M., Zhang, Y., Wang, H., Zhou, Y., Monteiro, M. J., Hao, L., Haughey, N. J., and Wang, J. (2026) UBQLN2 links proteotoxicity with lipid metabolism in neurodegeneration. Nat. Neurosci. 29, 782–795

37. Replogle, J. M., Marks, J. D., Fernandez, M. G., Yuan, H., Yu, D., Winters, E., Jawahar, V. M., Deshmukh, R., Sutanto, R., Kowal, I., Frankenfield, A., Shi, R., Carlomagno, Y., Jansen-West, K., Todd, T. W., Kopach, A., Ndayambaje, I. S., Qi, Y. A., Shantaraman, A., Pozo-Cabanell, I., Sheth, U., Yue, M., Duong, D., Ferguson, S. M., Bennett, D. A., Damme, M., Boeve, B. F., Day, G. S., Kellman, B., Skarnes, W. C., Petersen, R. C., Josephs, K. A., Graff-Radford, N. R., McDonough, J. A., Prudencio, M., Barmada, S. J., Zhang, Y., Hao, L., DeTure, M., Rawlinson, B., Engelberg-Cook, E., Casey, M. C., Perez, N., Dickson, D. W., Wingo, A., Liu, Y., Seyfried, N. T., Wingo, T. S., Mosalaganti, S., Petrucelli, L., and Ward, M. E. (2026) Neurodegeneration risk variants promote lysosomal TMEM106B fibril accumulation. bioRxiv,

38. Ryan, V. H., Lawton, S., Reyes, J. F., Hawrot, J., Frankenfield, A. M., Seddighi, S., Ramos, D. M., Epstein, J., Faghri, F., Johnson, N. L., Zou, J., Kampmann, M., Replogle, J., Qi, Y. A., Yuan, H., Johnson, K., Maric, D., Hao, L., Nalls, M. A., and Ward, M. E. (2025) Maintenance of neuronal TDP-43 expression requires axonal lysosome transport. Elife 14,

39. Frankenfield, A. M., Yang, K. L., Mazli, W. N. A. binti, Shih, J., Yu, F., Lo, E., Nesvizhskii, A. I., and Hao, L. (2025) Benchmarking SILAC Proteomics Workflows and Data Analysis Platforms. Molecular & Cellular Proteomics 24, 100980

40. Mathieson, T., Franken, H., Kosinski, J., Kurzawa, N., Zinn, N., Sweetman, G., Poeckel, D., Ratnu, V. S., Schramm, M., Becher, I., Steidel, M., Noh, K.-M., Bergamini, G., Beck, M., Bantscheff, M., and Savitski, M. M. (2018) Systematic analysis of protein turnover in primary cells. Nat. Commun. 9, 689

41. Dörrbaum, A. R., Kochen, L., Langer, J. D., and Schuman, E. M. (2018) Local and global influences on protein turnover in neurons and glia. Elife 7, e34202

42. Cavarischia-Rega, C., Sharma, K., Fitzgerald, J. C., and Macek, B. (2024) Proteome Dynamics in iPSC-Derived Human Dopaminergic Neurons. Molecular and Cellular Proteomics 23,

43. Yoshimori, T., Yamamoto, A., Moriyama, Y., Futai, M., and Tashiro, Y. (1991) Bafilomycin A1, a specific inhibitor of vacuolar-type H(+)-ATPase, inhibits acidification and protein degradation in lysosomes of cultured cells. Journal of Biological Chemistry 266, 17707–17712

44. Ritchie, M. E., Phipson, B., Wu, D., Hu, Y., Law, C. W., Shi, W., and Smyth, G. K. (2015) limma powers differential expression analyses for RNA-sequencing and microarray studies. Nucleic Acids Res. 43, e47–e47

45. Wang, C., Ward, M. E., Chen, R., Liu, K., Tracy, T. E., Chen, X., Xie, M., Sohn, P. D., Ludwig, C., Meyer-Franke, A., Karch, C. M., Ding, S., and Gan, L. (2017) Scalable Production of iPSC-Derived Human Neurons to Identify Tau-Lowering Compounds by High-Content Screening. Stem Cell Reports 9, 1221–1233

46. Fernandopulle, M. S., Prestil, R., Grunseich, C., Wang, C., Gan, L., and Ward, M. E. (2018) Transcription Factor–Mediated Differentiation of Human iPSCs into Neurons. Curr. Protoc. Cell Biol. 79, e51

47. Frankenfield, A., Ni, J., and Hao, L. (2022) Characterization of Neuronal Lysosome Interactome with Proximity Labeling Proteomics. Journal of Visualized Experiments,

48. Frankenfield, A. M., Ni, J., Ahmed, M., and Hao, L. (2022) Protein Contaminants Matter: Building Universal Protein Contaminant Libraries for DDA and DIA Proteomics. J. Proteome Res. 21, 2104–2113

49. Bruderer, R., Bernhardt, O. M., Gandhi, T., Miladinović, S. M., Cheng, L.-Y., Messner, S., Ehrenberger, T., Zanotelli, V., Butscheid, Y., Escher, C., Vitek, O., Rinner, O., and Reiter, L. (2015) Extending the Limits of Quantitative Proteome Profiling with Data-Independent Acquisition and Application to Acetaminophen-Treated Three-Dimensional Liver Microtissues. Molecular & Cellular Proteomics 14, 1400–1410

50. Salovska, B., Li, W., Di, Y., and Liu, Y. (2021) Boxcarmax: A high-selectivity data-independent acquisition mass spectrometry method for the analysis of protein turnover and complex samples. Anal. Chem. 93, 3103–3111

51. Cox, J., and Mann, M. (2012) 1D and 2D annotation enrichment: a statistical method integrating quantitative proteomics with complementary high-throughput data. BMC Bioinformatics 13, S12

52. Szklarczyk, D., Kirsch, R., Koutrouli, M., Nastou, K., Mehryary, F., Hachilif, R., Gable, A. L., Fang, T., Doncheva, N. T., Pyysalo, S., Bork, P., Jensen, L. J., and von Mering, C. (2023) The STRING database in 2023: protein–protein association networks and functional enrichment analyses for any sequenced genome of interest. Nucleic Acids Res. 51, D638–D646

53. Choi, M., Chang, C.-Y., Clough, T., Broudy, D., Killeen, T., MacLean, B., and Vitek, O. (2014) MSstats: an R package for statistical analysis of quantitative mass spectrometry-based proteomic experiments. Bioinformatics 30, 2524–2526

54. Schneider, C. A., Rasband, W. S., and Eliceiri, K. W. (2012) NIH Image to ImageJ: 25 years of image analysis. Nat. Methods 9, 671–675

55. Calzari, L., Barcella, M., Alari, V., Braga, D., Muñoz-Viana, R., Barlassina, C., Finelli, P., Gervasini, C., Barco, A., Russo, S., and Larizza, L. (2020) Transcriptome Analysis of iPSC-Derived Neurons from Rubinstein-Taybi Patients Reveals Deficits in Neuronal Differentiation. Mol. Neurobiol. 57, 3685–3701

56. Xu, X., Hua, X., Brown, K., Ren, X., and Zhang, Z. (2022) Mcm2 promotes stem cell differentiation via its ability to bind H3-H4. Elife 11, e80917

57. Fornasiero, E. F., Mandad, S., Wildhagen, H., Alevra, M., Rammner, B., Keihani, S., Opazo, F., Urban, I., Ischebeck, T., Sakib, M. S., Fard, M. K., Kirli, K., Centeno, T. P., Vidal, R. O., Rahman, R.-U., Benito, E., Fischer, A., Dennerlein, S., Rehling, P., Feussner, I., Bonn, S., Simons, M., Urlaub, H., and Rizzoli, S. O. (2018) Precisely measured protein lifetimes in the mouse brain reveal differences across tissues and subcellular fractions. Nat. Commun. 9, 4230

58. Jayapal, K. P., Sui, S., Philp, R. J., Kok, Y.-J., Yap, M. G. S., Griffin, T. J., and Hu, W.-S. (2010) Multitagging Proteomic Strategy to Estimate Protein Turnover Rates in Dynamic Systems. J. Proteome Res. 9, 2087–2097

59. Dou, Y., Qiu, D., Li, V., Wierzbińska, M. E., Keele, G. R., Liu, W., Yang, J., Paulo, J. A., Qi, L., and Zhang, T. (2025) Integral Synthesis and Clearance Analysis via DIA (ISDia) Reveals Coordinated Protein Dynamics Regulation during Endoplasmic Reticulum Stress. bioRxiv,

60. Toyama, B. H., Savas, J. N., Park, S. K., Harris, M. S., Ingolia, N. T., Yates, J. R., and Hetzer, M. W. (2013) Identification of Long-Lived Proteins Reveals Exceptional Stability of Essential Cellular Structures. Cell 154, 971–982

61. Maze, I., Wenderski, W., Noh, K.-M., Bagot, R. C., Tzavaras, N., Purushothaman, I., Elsässer, S. J., Guo, Y., Ionete, C., Hurd, Y. L., Tamminga, C. A., Halene, T., Farrelly, L., Soshnev, A. A., Wen, D., Rafii, S., Birtwistle, M. R., Akbarian, S., Buchholz, B. A., Blitzer, R. D., Nestler, E. J., Yuan, Z.-F., Garcia, B. A., Shen, L., Molina, H., and Allis, C. D. (2015) Critical Role of Histone Turnover in Neuronal Transcription and Plasticity. Neuron 87, 77–94

62. Cohen, L. D., Zuchman, R., Sorokina, O., Müller, A., Dieterich, D. C., Armstrong, J. D., Ziv, T., and Ziv, N. E. (2013) Metabolic Turnover of Synaptic Proteins: Kinetics, Interdependencies and Implications for Synaptic Maintenance. PLoS One 8, e63191

63. Heo, S., Diering, G. H., Na, C. H., Nirujogi, R. S., Bachman, J. L., Pandey, A., and Huganir, R. L. (2018) Identification of long-lived synaptic proteins by proteomic analysis of synaptosome protein turnover. Proceedings of the National Academy of Sciences 115,

64. Schanzenbächer, C. T., Sambandan, S., Langer, J. D., and Schuman, E. M. (2016) Nascent Proteome Remodeling following Homeostatic Scaling at Hippocampal Synapses. Neuron 92, 358–371

65. Zheng, X., Boyer, L., Jin, M., Mertens, J., Kim, Y., Ma, L., Ma, L., Hamm, M., Gage, F. H., and Hunter, T. (2016) Metabolic reprogramming during neuronal differentiation from aerobic glycolysis to neuronal oxidative phosphorylation. Elife 5, e13374

66. Mauvezin, C., and Neufeld, T. P. (2015) Bafilomycin A1 disrupts autophagic flux by inhibiting both V-ATPase-dependent acidification and Ca-P60A/SERCA-dependent autophagosome-lysosome fusion. Autophagy,

67. Papadopoulos, C., and Meyer, H. (2017) Detection and Clearance of Damaged Lysosomes by the Endo-Lysosomal Damage Response and Lysophagy. Current Biology 27, R1330–R1341

68. Turk, V., Stoka, V., Vasiljeva, O., Renko, M., Sun, T., Turk, B., and Turk, D. (2012) Cysteine cathepsins: From structure, function and regulation to new frontiers. Biochimica et Biophysica Acta (BBA) - Proteins and Proteomics 1824, 68–88

69. Martina, J. A., Diab, H. I., Lishu, L., Jeong-A, L., Patange, S., Raben, N., and Puertollano, R. (2014) The Nutrient-Responsive Transcription Factor TFE3 Promotes Autophagy, Lysosomal Biogenesis, and Clearance of Cellular Debris. Sci. Signal. 7,

70. Deutsch, E. W., Bandeira, N., Perez-Riverol, Y., Sharma, V., Carver, J. J., Mendoza, L., Kundu, D. J., Bandla, C., Kamatchinathan, S., Hewapathirana, S., Sun, Z., Kawano, S., Okuda, S., Connolly, B., MacLean, B., MacCoss, M. J., Chen, T., Zhu, Y., Ishihama, Y., and Vizcaíno, J. A. (2026) The ProteomeXchange consortium in 2026: making proteomics data FAIR. Nucleic Acids Res. 54, D459–D469

71. Perez-Riverol, Y., Bandla, C., Kundu, D. J., Kamatchinathan, S., Bai, J., Hewapathirana, S., John, N. S., Prakash, A., Walzer, M., Wang, S., and Vizcaíno, J. A. (2025) The PRIDE database at 20 years: 2025 update. Nucleic Acids Res. 53, D543–D553

