## Supplemental Information for "Neuronal Culture Media Shape Protein Abundance and Turnover in Human iPSC-Derived Neurons"

Ling Hao, Ph.D.

Associate Professor

Department of Chemistry & Biochemistry

University of Maryland

### Table of Contents


- **Supplemental Fig. S1.** Comparing four neuronal media with label-free proteomics.
- **Supplemental Fig. S2.** Pairwise differential abundance analysis among DMEM, Neurobasal, and DMEM+Neurobasal (1:1).
- **Supplemental Fig. S3.** Comparing four neuronal media with dynamic SILAC proteomics.
- **Supplemental Fig. S4.** GO enrichment analysis of changes in neuronal protein abundance following treatment with 50 nM BafA1.
- **Supplemental Fig. S5.** Combined differential analysis of neuronal protein synthesis (heavy) and degradation (light) following treatment with 50 nM BafA1 in BrainPhys and DMEM media.
- **Supplemental Table S1.** Molecular composition of BrainPhys, DMEM/F12, Neurobasal, and DMEM+Neurobasal (1:1) media.
- **Supplemental Table S2.** Label-free proteomics abundances from neurons cultured in four different neuronal media.
- **Supplemental Table S3.** Dynamic SILAC proteomics from neurons cultured in four different neuronal media.
- **Supplemental Table S4.** Label-free proteomics abundances from neurons with and without BafA1 treatment.
- **Supplemental Table S5.** Dynamic SILAC proteomics from neurons cultured in BrainPhys with and without BafA1 treatment. (A) light channel intensities; (B) heavy channel intensities; (C) heavy/light ratios**.**
- **Supplemental Table S6.** Dynamic SILAC proteomics from neurons cultured in DMEM with and without BafA1 treatment. (A) light channel intensities; (B) heavy channel intensities; (C) heavy/light ratios**.**

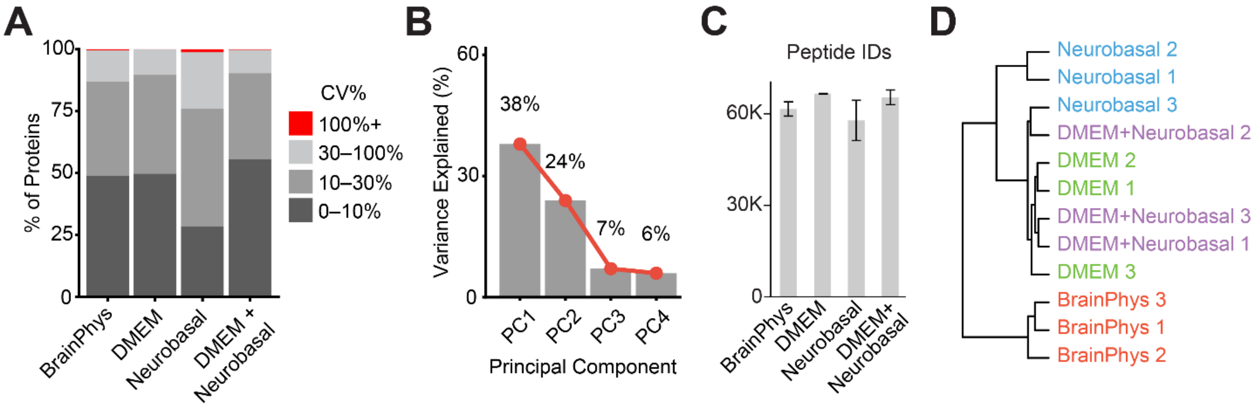


**Supplemental Fig. S1. Comparing four neuronal media with label-free proteomics.** (A) Distribution of coefficients of variation (CVs) in protein abundance across biological replicates for each medium. (B) PCA scree plot showing the variance explained by each principal component in Figure 1E. (C) Number of quantified peptides in each medium. (D) Hierarchical clustering of the neuronal protein abundance dataset across the four media.

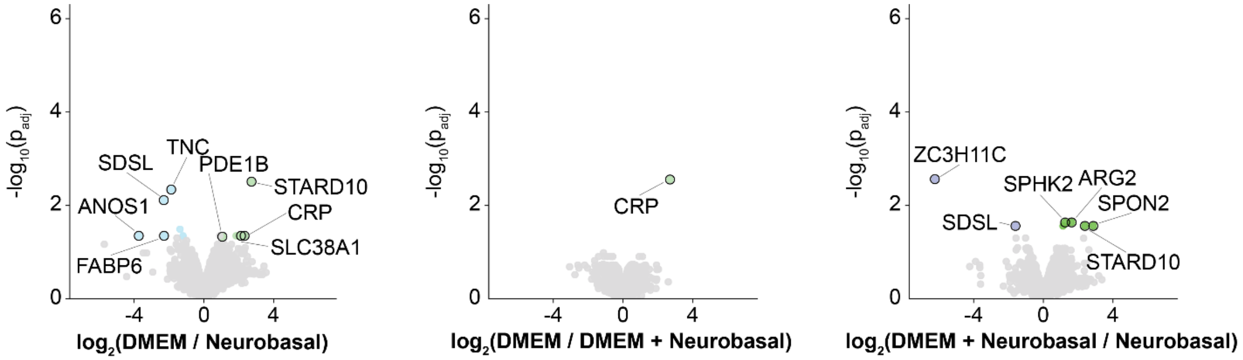


**Supplemental Fig. S2. Pairwise differential abundance analysis among DMEM, Neurobasal, and DMEM+Neurobasal (1:1).** Volcano plots comparing the three conventional media, showing minimal differences in relative protein abundance among them.

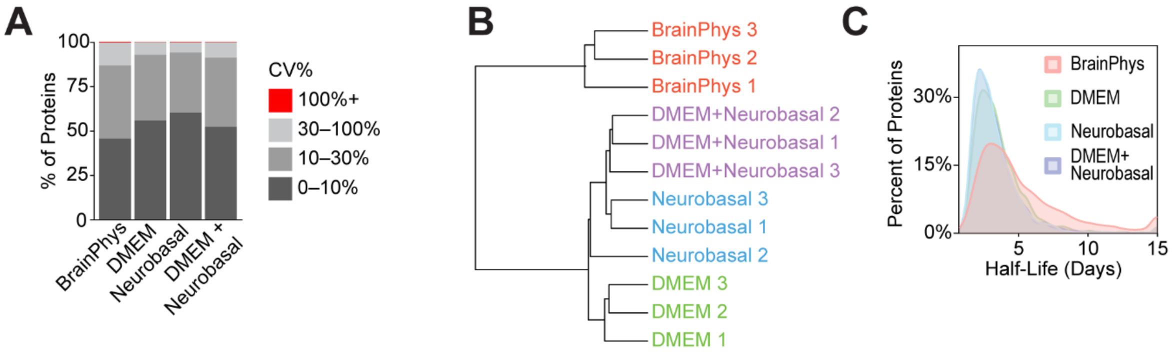


**Supplemental Fig. S3. Comparing four neuronal media with dynamic SILAC proteomics.** (A) Distribution of protein half-life CV categories for each medium. (B) Hierarchical clustering of protein half-lives across the four media. (C) Density plot of protein half-life distributions.

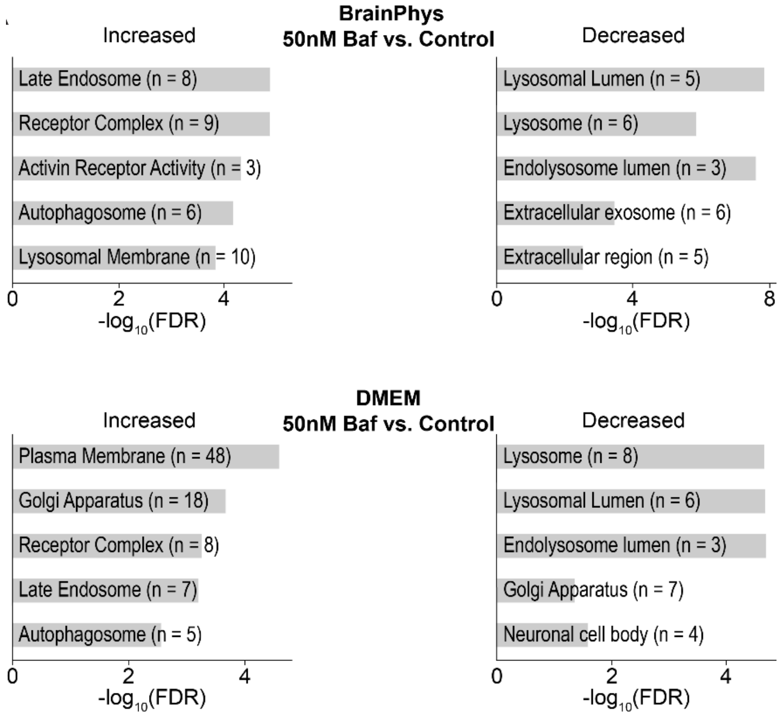


**Supplemental Fig. S4. GO enrichment analysis of changes in neuronal protein abundance following treatment with 50 nM BafA1.**

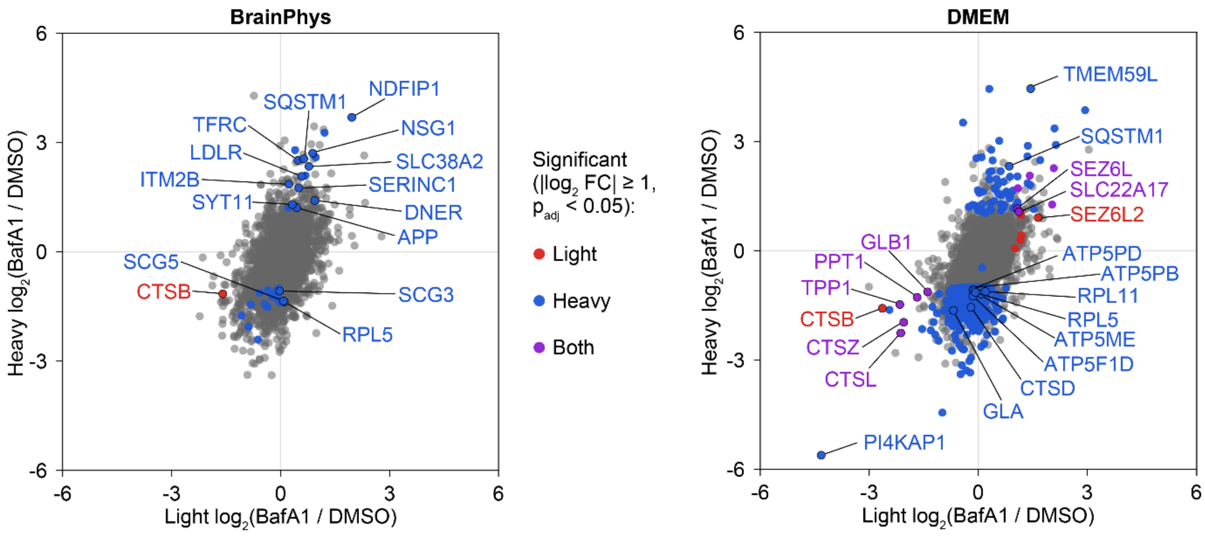


**Supplemental Fig. S5. Combined differential analysis of neuronal protein synthesis (heavy) and degradation (light) following treatment with 50 nM BafA1 in BrainPhys and DMEM media.**
